# Quantitative Analysis of Dopamine Neuron Subtypes Generated from Mouse Embryonic Stem Cells

**DOI:** 10.1101/093419

**Authors:** Yu-Ting L. Dingle, Katherine B. Xiong, Jason T. Machan, Kimberly A. Seymour, Debra Ellisor, Diane Hoffman-Kim, Mark Zervas

**Affiliations:** Department of Molecular Pharmacology, Physiology, and Biotechnology, Brown University, Providence, Rhode Island, USA; Center for Biomedical Engineering, Brown University, Providence, Rhode Island, USA; Department of Molecular Biology, Cell Biology, and Biochemistry, Brown University, Providence, Rhode Island, USA; Departments of Orthopedics and Surgery at Rhode Island Hospital and The Warren Alpert Medical School at Brown University, Providence, Rhode Island, USA; Brown Institute for Brain Science, Providence, Rhode Island, USA; School of Engineering, Brown University, Providence, Rhode Island, USA

## Abstract

Dopamine (DA) neuron subtypes modulate specific physiological functions and are involved in distinct neurological disorders. Embryonic stem cell (ESC) derived DA neurons have the potential to aid in the study of disease mechanisms, drug discovery, and possibly cell replacement therapies. DA neurons can be generated from ESCs in vitro, but the subtypes of ESC-derived DA neurons have not been investigated in detail despite the diversity of DA neurons observed in vivo. Due to cell culture heterogeneity, sampling methods applied to ESC-derived cultures can be ambiguous and potentially biased. Therefore, we developed a quantification method to capture the depth of DA neuron production in vitro by estimating the error associated with systematic random sampling. Using this method, we quantified calbindin+ and calretinin+ subtypes of DA neurons generated from mouse ESCs. We found a higher production of the calbindin+ subtype (11−27%) compared to the calretinin+ subtype (2-13%) of DA neuron; in addition, DA neurons expressing neither subtype marker were also generated. We then examined whether exogenous sonic hedgehog (SHH) and fibroblast growth factor 8 (FGF8) affected subtype generation. Our results demonstrate that exogenous SHH and FGF8 did not alter DA neuron subtype generation in vitro. These findings suggest that a deeper understanding DA neuron derivation inclusive of mechanisms that govern the in vitro subtype specification of ESC-derived DA neurons is required.

**Note:** All research was planned and conducted while members were at Brown University

**Research funding:** NIH/NCRR/NIGMS RI Hospital COBRE Center for Stem Cell Biology (8P20GM103468-04) (MZ) Brown Institute for Brain Science Pilot Grant (4-63662) (MZ/DHK)

## INTRODUCTION

Dopamine (DA) neurons in the midbrain are a diverse population and consist of subtypes with distinct anatomical and molecular identities [1]. When defined by anatomical region, the major groups are the substantia nigra (SN) or the A9 group and the ventral tegmental area (VTA) or the A10 group [2]. SN DA neurons, which innervate the dorsal striatum, are essential for initiating movement and controlling postural reflexes [1, 3–5]. VTA DA neurons, which project to the ventral striatum, limbic, and cortical areas, are involved in cognition, reward, and emotional behavior [2, 6, 7]. Degeneration of SN DA neurons leads to the loss of motor function in Parkinson's disease [8], while abnormal activity of VTA DA neurons may underlie schizophrenia, addiction, and attention deficit hyperactivity disorder [1, 3, 7, 9, 10].

The heterogeneous molecular identity of DA neurons adds further complexity to their subtype characterization. Calbindin (CALB), a calcium-binding protein, is enriched in the majority of VTA DA neurons and in a subset of SN DA neurons [11–16]. A recent study suggested that CALB buffers DA vesicle release in healthy VTA neurons [17]. The expression of CALB may protect DA neurons from calcium cytotoxicity-induced degeneration as VTA DA neurons are less affected than SN DA neurons in Parkinson’s disease [12, 18, 19]. Calretinin (CALR), another calcium-binding protein, is expressed in DA neurons scattered in the VTA and SN [14–16]. The physiological function of CALR is not well understood, but it has been suggested that CALR may have similar neuroprotective properties to CALB [20, 21]. A third protein, G-protein-regulated inward-rectifier potassium channel 2 (GIRK2), is expressed in nearly all SN DA neurons, which undergo the greatest degree of degeneration in Parkinson’s disease [11, 22, 23]. The expression of CALB, CALR, and GIRK2 is not mutually exclusive and gives rise to diverse subtypes of DA neurons [14–16, 24, 25].

Because of their clinical relevance, the generation of DA neurons *in vitro* offers opportunities to study disease mechanism, progression and therapeutics, and holds potential for cell replacement therapy [26–29]. A well established method of deriving DA neurons from embryonic stem cells (ESCs) and induced pluripotent stem cells (iPSCs) *in vitro* is a 5-stage differentiation method first described by Lee et al. [30]. Each stage utilizes a defined culture substrate and media and cells become lineage-restricted as they pass through each stage. Differentiated neurons are observed at the end of the protocol with DA neurons accounting for a small percentage of all neurons. The DA neuron population generated using this 5-stage method is not homogeneous [31–33] and the subtype profile has not been quantitatively analyzed.

The efficiency of DA neuron production with the 5-stage method may be enhanced by exposing neural progenitors at the fourth stage to exogenous sonic hedgehog (SHH) and fibroblast growth factor 8 (FGF8). SHH and FGF8 are two morphogens required for inducing DA neuron fate in the midbrain during embryonic development *in vivo* [34]. At embryonic day 8.5 (E8.5) SHH is expressed along the ventral midline and FGF8 in the isthmus at the mid/hindbrain boundary [35]. The intersection of these two morphogens is where DA neuron progenitors appear during embryogenesis [34]. A SHH gradient has been shown to be involved with motor neuron subtype specification in the spinal cord [36], but it is unknown whether soluble morphogens play a role in the subtype specification of DA neurons *in vitro*.

In this study, we established a sampling approach to quantify DA neuron subtypes in heterogeneous cultures. We evaluated the generation of DA neuron subtypes from a baseline 5-stage culture, in which no exogenous SHH and FGF8 were added. Among the DA neurons that were generated we detected the expression of CALB and CALR, but not GIRK2. We then conducted quantitative analysis of the CALB+ and CALR+ subtypes of DA neurons. We further investigated whether the distribution of CALB+ and CALR+ subtypes could be manipulated by the addition of SHH and FGF8 at different stages. We discuss these findings in the context of subtype specification of DA neurons *in vitro*.

## MATERIALS AND METHODS

### Mouse embryonic stem cell culture

Mitomycin c-inactivated mouse embryonic fibroblasts (iMEFs), obtained from Brown University’s Transgenic Facility, were cultured for 24 hrs in MEF media [Dulbecco’s Modified Eagle Medium (DMEM, Invitrogen, 11965), 10% fetal bovine serum (FBS, Gemini, 100-106) and 1% Penicillin-Streptomycin-Glutamine (Pen/Strep/Glut, Invitrogen, 10378)] in 10 cm tissue culture dishes (BD, 353003) coated with 0.1% gelatin (Sigma, G2500). D3 ESCs (ATCC, CRL-11632) and R1 ESCs (ATCC, SCRC-1011) were each cultured on confluent iMEF monolayers at 2.5-3x10^6^ cells/dish in ES media [Knock-out DMEM (Invitrogen, 10829), 15% FBS, 100 μM MEM nonessential amino acids (Invitrogen, 11140), 1% Pen/Strep/Glut, and 55 μM 2-mercaptoethanol (Invitrogen, 21985)] supplemented with 1400 U/mL ESGRO leukemia inhibitory factor (LIF, Millipore, ESG1107). ESCs were cultured for 1-2 days before colonies contacted each other and were dissociated using 0.05% trypsin (Invitrogen, 25300). The ESCs were passaged onto a new iMEF monolayer, cryopreserved in ES media with an additional 10% FBS and 10% DMSO (Sigma, D2650), or used for DA neuron differentiation (see below). Passage number of D3 ESCs was not provided by ATCC, and therefore we designated passage 1 (P1) as the purchased vial.

### DA neuron differentiation

We used a 5-stage protocol, with a baseline culture that was slightly modified from the 5-stage method first described by Lee et al. in 2000 [30] and commercialized by R&D Systems (Human/ Mouse DA Neuron Differentiation Kit, R&D Systems, SC001b). The five stages of baseline culture conditions were as follows (illustrated in **Figure 1** and shown in **Supplementary Figure 1**). *Stage I* (ESC expansion): ESCs were cultured on gelatin-coated tissue culture dish in ESC media with LIF for 1-2 days before colonies contacted each other. *Stage II* (embryoid body (EB) formation): ESC colonies were dissociated with 0.05% trypsin. Monodispersed ESCs were cultured in non-adherent bacterial culture dishes (BD, 351029) at 2x10^6^ cells/dish for 3-4 days in ES media without ESGRO to form EBs. *Stage III* (neural precursor (NP) selection): EBs were transferred to tissue culture dishes in ES media. After 24 hrs, the media was replaced with ITSFn media [DMEM/F12 (Invitrogen,11320), 1X insulin-transferrin-selenium supplement (ITS, Invitrogen, ITS-G), 50 µg/mL fibronectin (Fn, R&D Systems, 1030-FN), and 1% Pen/Strep (Invitrogen, 15140)]. The cells were cultured in ITSFn media for 6 days and then dissociated with 0.05% trypsin. *Stage IV* (NP expansion): Monodispersed Nestin+ NPs were cultured at 3x10^5^ cells/well on 12-mm glass coverslips coated with 15 µg/mL poly-L-ornithine (PLO, Sigma, P4957) and 1 µg/mL Fn in 24-well plates in N2aa media [DMEM/F12, N-2 Max supplement (R&D, AR009), 1% Pen/Strep, and 200µM ascorbic acid (Sigma, A4403)] supplemented with 10 ng/mL basic fibroblast factor (bFGF, R&D, 233-FB-025/CF). Cells were cultured for 4 days. *Stage V* (Differentiation): Media was replaced with N2aa media without bFGF. Cells were cultured in this differentiation media for either 10 days for D3 or 15 days for R1.

**Figure 1.**
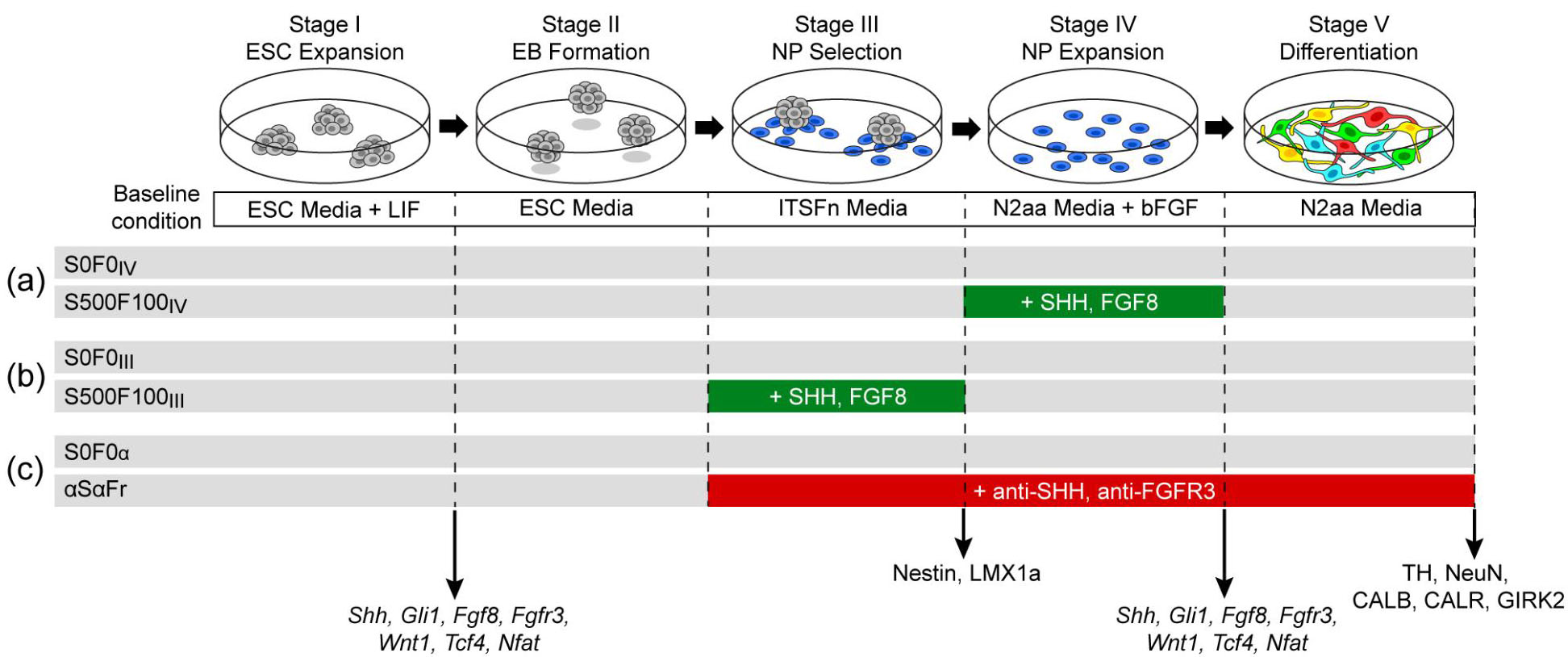
Summary of culture conditions. A modified version of the 5-stage protocol was used inthis study. Baseline control conditions (S0F0_IV_, S0F0_III_, and S0F0_α_): Undifferentiated ESCs were expanded in ESC media with LIF (Stage I, ESC Expansion). ESCs were cultured as EBs in suspension in ESC media without LIF (Stage II, EB Formation). NPs (blue) were selected in ITSFn media (Stage III, NP Selection). NPs were expanded in N2aa media with bFGF (10 ng/mL) (Stage IV, NP Expansion). Cells differentiated after growth factor withdrawal (Stage V, Differentiation). Experimental conditions: **(a)** SHH (500 ng/mL) and FGF8 (100 ng/mL) were added during Stage IV (S500F100_IV_). The paired control (S0F0_IV_) was conducted in parallel. **(b)** SHH (500 ng/mL) and FGF8 (100 ng/mL) were added during Stage III (S500F100III). The paired control (S0F0_III_) was conducted in parallel. **(c)** Anti-SHH (10 µg/mL) and anti-FGFR3 (10 µg/mL) antibodies were added at the start of Stage III until the end of the experiment (αSαFr). The paired control (S0F0_α_) was conducted in parallel. Differentiated cells at the end of Stage V were immunolabeled for TH (DA neurons), NeuN (neurons), and subtype markers CALB, CALR, and GIRK2. NPs at the end of Stage III were immunolabeled for Nestin (NP) and LMX1a (transcription factor associated with DA neuron progenitors). Undifferentiated ESCs and cells at the end of Stage IV were analyzed for the mRNA expression of SHH and FGF8 signaling components, namely *Shh, Gli1, Fgf8, Fgfr3*, Wnt1, Tcf4, and Nfat.

In the first experiment, passage 3 (P3) of D3 ESCs and P21 of R1 ESCs were used (**Figure 1a**, S500F100_IV_,). SHH (500 ng/mL, R&D, 461-SH-025/CF) and FGF8 (100 ng/mL, R&D, 423-F8-025/CF) were added to N2aa media and refreshed upon media exchange during NP expansion (Stage IV). In the second experiment, P5 of D3 ESCs was used, and SHH and FGF8 were added to the ITSFn media during NP selection (Stage III) and refreshed upon media exchange (**Figure 1b**, S500F100III). In the third experiment, P5 of D3 ESCs was used (**Figure 1c**, αSαFr). Anti-SHH (10 µg/mL, R&D, MAB4641) and anti-FGFR3 (10 µg/mL, R&D, MAB710) antibodies were added at the beginning of Stage III to the end of Stage V. Antibodies were refreshed upon media change.

A common problem with ESC culture is the variation among sub-clones [37]. To remove confounding variables, all experiments described in this study were conducted in parallel with a paired control group (S0F0_IV_, S0F0_III_, and S0F0_α_) using the baseline culture protocol. The experimental and control groups started with the same passage and subclone of ESCs. To evaluate the expression of tyrosine hydroxylase (TH), NeuN, CALB, CALR, and GIRK2 in the differentiated culture, cells were fixed with 4% paraformaldehyde at the end of Stage V. To evaluate the expression of Nestin and LMX1a in NPs, cells at the end of Stage III were trypsinized and re-plated on PLO/Fn-coated glass coverslips in ITSFn media for 24 hrs and then fixed.

### Immunocytochemistry

Immunocytochemistry was carried out using standard protocols [14]. Primary antibodies were diluted in working solution [phosphate buffered saline (PBS), 10% normal donkey serum and 0.2% Triton X-100] with the following dilutions: mouse anti-nestin (Millipore, MAB353, 1:200), rabbit anti-LMX1a (Millipore, AB10533, 1:200), mouse anti-Tuj-1 (R&D, MAB1195, 1:1000), rabbit anti-TH (Millipore, AB152, 1:1000), sheep anti-TH (Millipore, AB1542, 1:1000), rabbit anti-calbindin (SWANT, 1:5000), goat anti-calretinin (Millipore, AB1550, 1:5000). Mouse anti-NeuN antibody (Millipore, MAB377, 1:100) was diluted in PBS. Primary antibody incubation was done at 4°C overnight, followed by three washes with PBT [0.2% Triton-X in PBS]. Appropriately matched secondary antibodies were diluted in working solution (all at 1:500): Alexa 488 donkey anti-sheep (Invitrogen, A11015), Alexa 488 donkey anti-rabbit (Invitrogen, A21206), Alexa 555 donkey anti-mouse (Invitrogen, A31570), Alexa 555 donkey anti-rabbit (Invitrogen, A31572), and Alexa 555 donkey anti-goat (Invitrogen, A21432). Secondary antibody incubation was done at room temperature for 1 hour, followed by three PBT washes. Nuclei were counter-stained with Hoechst or DAPI. All experiments were performed with n ≥ 3 coverslips. 20X z-stack images with 2-µm z-steps were acquired with Volocity 5.2 software and a Leica fluorescence microscope (DM6000B) or a Nikon Eclipse (TS2000) fluorescence microscope. All images were pseudo-colored during acquisition. Raw images were used for quantification. Images shown in the figures are z-stacks (typical thickness 20-40 µm) reconstructed using the Extended Focus function in Volocity to show cells in focus from multiple z-planes, and brightness and contrast were adjusted using the level function in Photoshop for better visualization of markers.

### Quantification

To quantify neurons that expressed markers of interest, z-stack raw images were opened in Volocity with one single z-plane shown at a time. The green channel with TH+ labeling was uncloaked while the other channels were cloaked. TH+ cells in each field of view (FOV) were tagged using the point tool. TH+ cells were examined through all the z-planes to ensure that each object was counted only once. The red channel (CALB, CALR, or NeuN labeling) was then uncloaked to determine double-positive cells in each FOV. The percent of DA neurons was calculated as the number of TH+ cells divided by NeuN+ cells. We used NeuN because it is restricted to neuronal nuclei and facilitates more accurate counting then using TUJ-1, which labels neuronal soma and processes. The percent of CALB+ DA neurons was calculated as the number of TH+CALB+ cells divided by TH+ cells, and the percent of CALR+ DA neurons was calculated as the number of TH+CALR+ cells divided by TH+ cells.

### Sampling size selection by Monte Carlo simulation

The Monte Carlo error model was used to estimate error and help determine our sub-sampling method. The model was based on large coverage sampling of DA neuron-containing cultures by imaging eighty non-overlapping fields of view (operationally, we define multiple fields as FOVs) with a 20X objective; this was applied to two separate culture samples. The number of TH+ neurons in each FOV was manually counted as described above. A Matlab simulation was developed to randomly select subsets of N numbers of FOVs and calculate the average number of TH+ neurons based on the actual counts from the subset of FOVs. The simulation was executed 1000 times for all subset sizes. The standard errors were calculated from the standard deviations of the 1000 averages for each sampling size.

### Quantitative Real Time PCR (qRT-PCR)

Three experimental groups including undifferentiated ESCs and NPs derived from S0F0_IV_ and S500F100_IV_ conditions were harvested for RNA extraction. For each of the groups, multiple wells (n > 2) were collected to generate biological replicates. RNA was extracted using Trizol (Invitrogen, 15596-026) and purified using a RNeasy Mini Kit (Qiagen, 74104) according to manufacturers' instructions. RNA was quantified using Nanodrop ND-1000. A 0.5 µg RNA template was used to synthesize cDNA using iScript cDNA Synthesis Kit (BioRad, 170-8890) according to manufacturer's instruction. The cDNA from each biological replicate was run in three separate qRT-PCR reactions (in triplicate) for each primer set. Quantitative RT-PCR (qRT-PCR) was performed with SYBR Green (Applied Biosystems, 4367650) using ABI 7900 Fast Sequence Detection Instrument. Expression of *Shh*, *Gli1*, *Fgf8*, *Fgfr3, Wnt1, Tcf4, and Nfat* were analyzed and normalized to 18S. Results are reported as mean cycle threshold, standard deviation, and significance determined by two-tailed p value. Cycle times were graphed using a log(2) scale to match cycle times. See S1 Table for primer sequences.

### Statistical Method

Thirty FOVs were analyzed for each marker combination in each culture sample. Statistical analyses of TH+/mm^2^, NeuN+/mm^2^, TH+/NeuN+, TH+CALB+/TH+, and TH+CALR+/TH+ neurons were performed in SAS software and Generalized Estimating Equations were used to test all hypotheses. FOVs within each coverslip were modeled to have correlated residual error. For analyses comparing the percent of one type of cell within a larger pool of cells (e.g. TH+CALB+/ TH+, TH+CALR+/TH+, and TH+/NeuN+) a binomial distribution was used, which describes the probability distribution of the number of successes (e.g. TH+CALB+) in a sequence of trials (e.g. TH+). Binomial distribution appropriately constrains values between 0-100%, models the way the variability of percentages skewed above and below 50%, and scales the variability by the size of the denominator. For analyses comparing the number of TH+ neurons per area (e.g. TH+/mm^2^) a Poisson distribution was used, which describes the number of events (e.g. TH+) one would observe in a fixed event rate (events per mm^2^). Unlike binomial distribution, Poisson distribution appropriately constrains values only to ≥ 0.

Once the appropriate distribution models were fit for the dependent variable (TH+/mm^2^, TH+/NeuN+, TH+CALB+/TH+, and TH+CALR+/TH+), individual orthogonal linear estimates were constructed within the models to test specified hypotheses. Families were defined within a dependent variable as comparisons between experimental conditions and their paired controls. Familywise alpha was maintained at 0.05 using the Holm test to create adjusted p-values, and statistical significance was defined as adjusted p < 0.05. Comparisons among controls were also conducted to assess the variability among ESC lines and passages. Finally, classical sandwich estimation was used to adjust the final model parameters based on how closely the parameters fit the distribution of the data. Means and 95% confidence intervals (CI) are reported.

## RESULTS

### Quantification using Monte Carlo simulation

We generated DA neurons from ESCs using a modified version of the 5-stage method (See Methods, **Figure 1**). Immunolabeling for TH revealed that the *in vitro* distribution of TH+ (DA) neurons generated from ESCs was highly heterogeneous (**Figure 2a** and **Supplemental Figure 2**). To quantitatively evaluate our cultures in an unbiased manner we selected a sampling size to ensure accuracy while taking into account sampling error and efficiency. As illustrated in **Figure 2b**, we sampled 80 evenly spaced FOVs positioned in the center of the coverslip with a 1 FOV-space between sampled FOVs. We made the assumption that 80 FOVs yielded sufficient coverage to represent the entire culture. We manually counted the number of TH+ neurons in each FOV and calculated the average from the 80 FOV counts (Average TH+_80_) (**Figure 2c**). We did the same for TH+CALB+ neurons from the 80 FOV counts (Average TH+CALB+_80_) (**Figure 2d**). We then used a Monte Carlo error model to assess the error associated with randomly sampling smaller subsets from the 80 FOVs (**Figure 2e**). Specifically, we defined sample size N, where 0 < N < 80. A set of N numbers of FOVs was randomly selected from the 80 FOVs, and the average TH+ counts from this subset was calculated (Average TH+_N_). This process was executed 1000 times for each sample size N. The 1000 Averages TH+_N_ deviated from Average TH+_80_ to varying extents (**Figure 2f**). The standard deviation of the 1000 averages was calculated, which by definition is the standard error of the sample (**Figure 2h**, green line). As expected, it was found that with larger sampling size there was a smaller resulting error. For example, the errors were ±27.43% for 10 FOV, ±17.04% for 20 FOV, ±13.05% for 30 FOV, and ±9.75% for 40 FOV. Taking into consideration the error and the efficiency of performing the counts, we concluded that 30 FOVs per sample provided sufficient representation of the cultures and an acceptable amount of error. We repeated this process for TH+CALB+ neurons (**Figures 2d, g, h**) and reached a similar conclusion. Subsequently, all cultures described in this study were analyzed by imaging 30 randomly selected FOVs.

**Figure 2.**
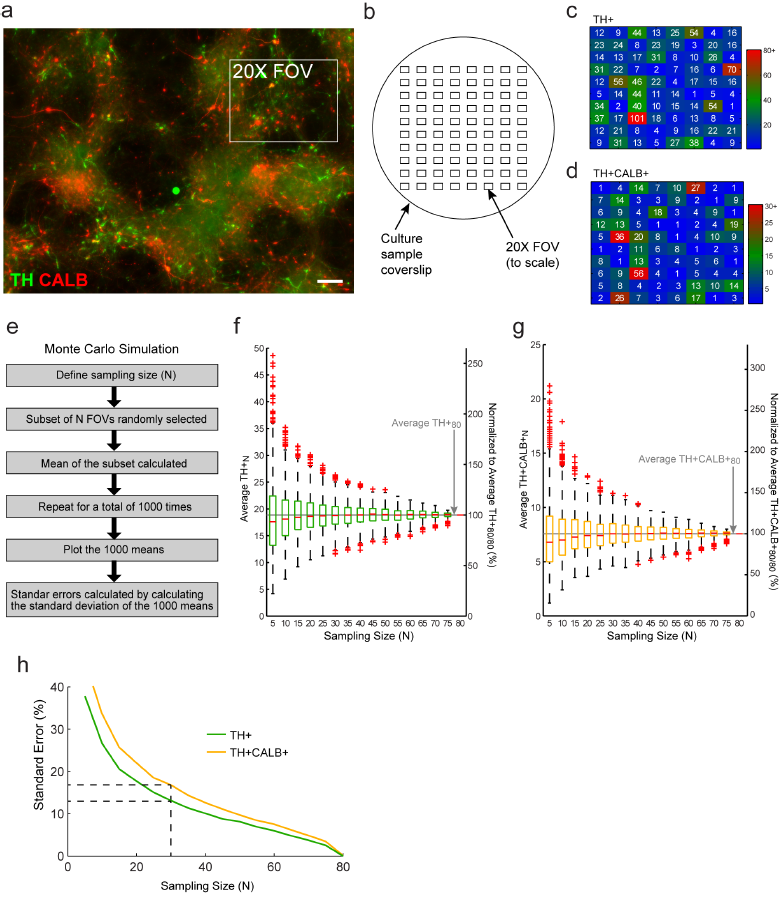
The spatial distribution of differentiated neurons was highly heterogeneous. To determine a sufficiently accurate quantitative estimation, we evaluated the error associated with sampling size. **(a)** A stitched fluorescence image shows the distribution of TH+ (green) and CALB+ (red) neurons. The box indicates a 20X field of (FOV). Scale bar, 100 µm. **(b)** A high coverage of cultures grown on coverslips was obtained by the representative sampling of 80 FOVs. The drawing in (b) shows the size and spacing of the 80 FOVs relative to the coverslip (diameter 10 mm). Each FOV was 1 FOV-distance apart from the next one. **(c, d)** The numbers of TH+ neurons **(c)** and TH+CALB+ neurons **(d)** in each FOV were manually counted. The numbers are the actual counts in corresponding FOVs. The averages from all 80 FOVs were calculated (Average TH+80, Average TH+CALB+80). **(e)** A Monte Carlo simulation in Matlab was used to randomly choose a subset of the FOVs with a defined sampling size (N = number of FOVs) among the 80. The average was calculated from the subset (Average TH+N, Average TH+CALB+N). This process was repeated to generate 1000 averages for each sampling size N **(f,g**). Representative box plots of the 1000 Average TH+N **(f)** and Average TH+CALB+N **(g)** from a simulation run of each sampling size N. Red dashed lines show the medium values for each simulation and box boundaries show the 25th and 75th percentiles. Red crosses indicate outliers. Average TH+N and Average TH+CALB+N are shown as the gray lines. **(h)** Representative graph of the standard error (%) vs. sampling size (N) from a simulation. Dashed lines indicate the chosen 30 FOVs for our study and the corresponding standard errors for TH+ and TH+CALB+.

### 5-stage method generated more CALB+ than CALR+ subtype in D3-derived DA neurons

We assessed the generation of DA neuron subtypes by double immunolabeling for TH and CALB, TH and CALR, or TH and GIRK2 following a 5-stage culture paradigm in the absence of exogenous SHH and FGF8. We observed qualitatively more TH+CALB+ DA neurons versus TH+CALR+ DA neurons which was verified by quantitation (**Table 1** and **Figure 3a, c**). We also observed many TH-CALB+ neurons and a smaller number of TH-CALR+ neurons, presumably interneurons, which were not quantified in our study. We did not detect TH+GIRK2+ neurons at the end of the 10-day differentiation period. GIRK2+ expression was observed when we extended differentiation to 25 days (**Supplemental Figure 3**). For the purpose of this study, we restricted our quantitative analysis to the standard differentiation duration typically cited in the literature [30]. The percentage of TH+DA neuron subtypes generated with the S0F0_IV_ baseline culture condition was 26.24% for the CALB+ subtype, which was significantly higher than 13.14% for the TH+CALR+ subtype (p < 0.0001) (**Table 1** and **Figure 3e**).

**Table 1.** Quantification and statistical comparisons of CALB+ and CALR+ subtypes. The percentage of CALB+ and CALR+ subtypes of DA neurons with adjusted p values. CI, confidence interval (95%).

| **Table 1. Quantification and statistical comparisons of CALB+ and CALR+ subtypes** |  |  |  |  |  |  |  |  |
| --- | --- | --- | --- | --- | --- | --- | --- | --- |
| Conditions |  | %TH+CALB+/TH+ |  |  | %TH+CALR+/TH+ |  |  | CALB+ vs CALR+ |
|  |  | Mean | Lower CI | Upper CI | Mean | Lower CI | Upper CI |  |
| D3 | S0F0<sub>IV</sub> | 26.24 | 24.63 | 27.92 | 13.14 | 12.04 | 14.31 | $p < 0.0001$ |
| | S500F100<sub>IV</sub> | 27.25 | 24.44 | 30.26 | 10.95 | 10.33 | 11.61 | $p < 0.0001$ |
| | | $p = 1.000$ | | | $p = 0.010$ | | | |
| | S0F0<sub>III</sub> | 12.31 | 11.89 | 12.75 | 8.53 | 6.35 | 11.37 | $p = 0.0003$ |
| | S500F100<sub>III</sub> | 11.01 | 10.33 | 11.73 | 8.81 | 8.03 | 9.66 | $p = 0.0142$ |
| | | $p = 0.025$ | | | $p = 1.000$ | | | |
| | S0F0<sub>α</sub> | 12.49 | 10.11 | 15.33 | 9.04 | 8.17 | 9.99 | $p < 0.0001$ |
| | αSaFr | 12.43 | 11.70 | 13.20 | 8.21 | 7.81 | 8.63 | $p = 0.0084$ |
| | | $p = 1.000$ | | | $p = 0.467$ | | | |
| R1 | S0F0<sub>IV</sub> | 10.92 | 9.43 | 12.63 | 2.41 | 1.83 | 3.16 | $p < 0.0001$ |
| | S500F100<sub>IV</sub> | 10.87 | 9.59 | 12.29 | 3.49 | 2.95 | 4.11 | $p < 0.0001$ |
| | | $p = 1.000$ | | | $p = 0.144$ | | | |

### The role of exogenous SHH and FGF8 during Stage IV on DA neuron subtype generation from D3 ESCs

The addition of SHH and FGF8 during NP expansion (Stage IV), but not prior or after, has been reported to enhance DA neuron production [30]. To study if exogenous SHH and FGF8 shape DA neuron subtype generation, we supplemented culture media with exogenous SHH (500 ng/mL) and FGF8 (100 ng/mL) during Stage IV (500F100_IV_,) in parallel to the S0F0_IV_ baseline culture (**Figure 3**). The S500F100_IV_ conditions yielded 27.25% CALB+ DA neurons (**Figure 3a, b, e**), which was significantly higher than 10.95% of the CALR+ DA neuron subtype (**Figure 3c, d, e**) (p < 0.0001) (**Table 1**). These findings were similar between the S0F0_IV_ and S500F100_IV_ groups (**Table 1**). The comparison of CALB+ subtype across the S0F0_IV_ and S500F100_IV_ groups revealed no difference (p = 1.000) while the comparison of CALR+ subtype across the S0F0_IV_ and S500F100_IV_ groups revealed a 2.19% difference (p = 0.010).

**Figure 3.**
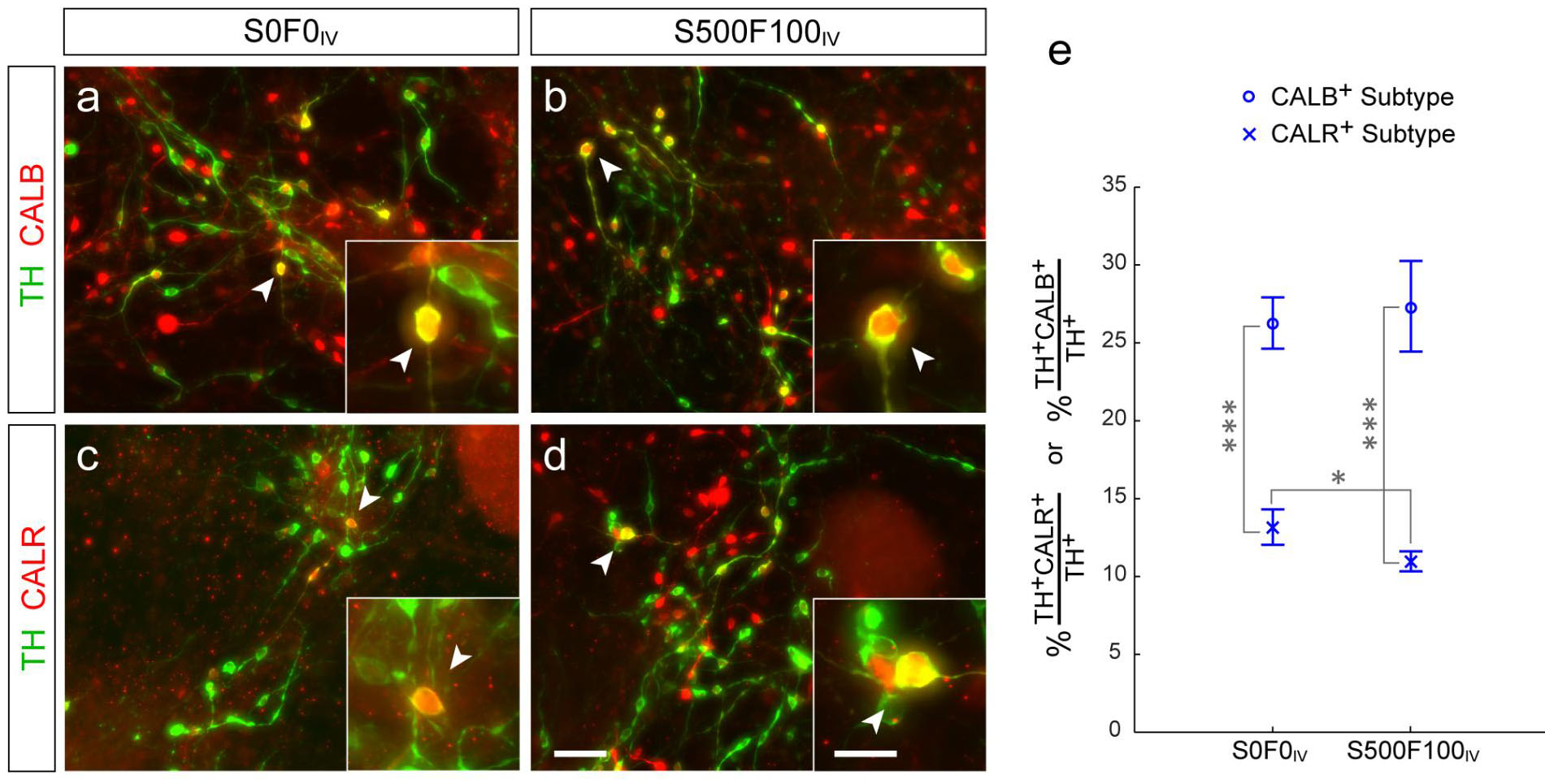
Quantification of CALB+ and CALR+ DA neuron subtypes in S0F0_IV_ vs S500F100_IV_. D3 ESCs cultured in S0F0_IV_ baseline or S500F100_IV_ conditions in which exogenous SHH (500 ng/ml) and FGF8 (100 ng/ml) were added during NP expansion (Stage IV). **(a, b)** Images of TH (green) and CALB (red) immunofluorescent labeling in S0F0_IV_ **(a)** and S500F100_IV_ **(b)**. **(c, d)** Images of TH (green) and CALR (red) immunofluorescent labeling in S0F0_IV_ **(c)** and S500F100_IV_ **(d)**. Arrowheads indicate examples of neurons that are TH+CALB+ or TH+CALR+; high magnification images of the same regions are shown in the insets. **(e)** Quantification of CALB+ and CALR+ DA neuron subtypes revealed a higher percent of CALB+ than CALR+ subtype in both S0F0_IV_ and S500F100_IV_ conditions. The addition of SHH and FGF8 resulted in a small, but statistically significant reduction in CALR+ subtype. Means and 95% CI are shown with values provided in Table 1. * p = 0.010, *** p < 0.0001. Scale bar (a - d), 50 μm; insets, 20 μm.

### Exogenous SHH and FGF8 during NP expansion did not influence DA neuron production

To validate the effect of exogenous morphogens on total DA neuron production we asked whether exogenous SHH and FGF8 in the S500F100IV condition altered the production of TH+ neurons from D3 ESCs compared to the S0F0IV condition. The total TH+ neuron production in S0F0IV (123.12 TH+/mm2) was not significantly different from S500F100_IV_ (98.34 TH+/mm2, p = 0.406) (**Table 2** and **Figure 4**). The total neuron production in S0F0_IV_ (2,110 NeuN+/mm2) was not significantly different from S500F100IV (1,867 NeuN+/mm2, p = 0.877). The percent of TH+ neurons within the total NeuN+ neuronal population in S0F0IV (6.18%) was also not significantly different from S500F100IV (6.49%, p = 1.000). This data shows that exogenous SHH and FGF8 did not significantly affect total neuron or DA neuron production derived from D3 ESCs.

**Table 2.**
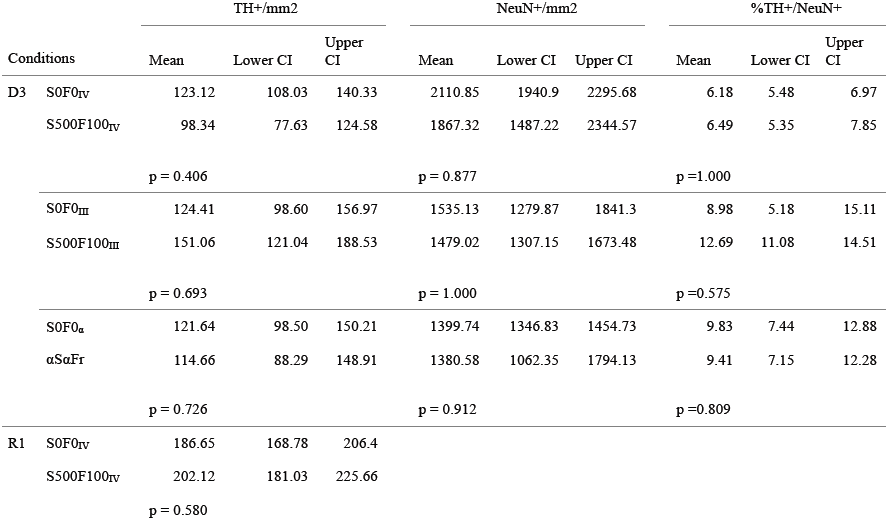
Quantification and statistical comparisons of total neurons and DA neurons. DA neurons (TH+), total neurons (NeuN+), percent DA neurons versus total neurons, and adjusted p values. CI, confidence interval (95%).

**Figure 4.**
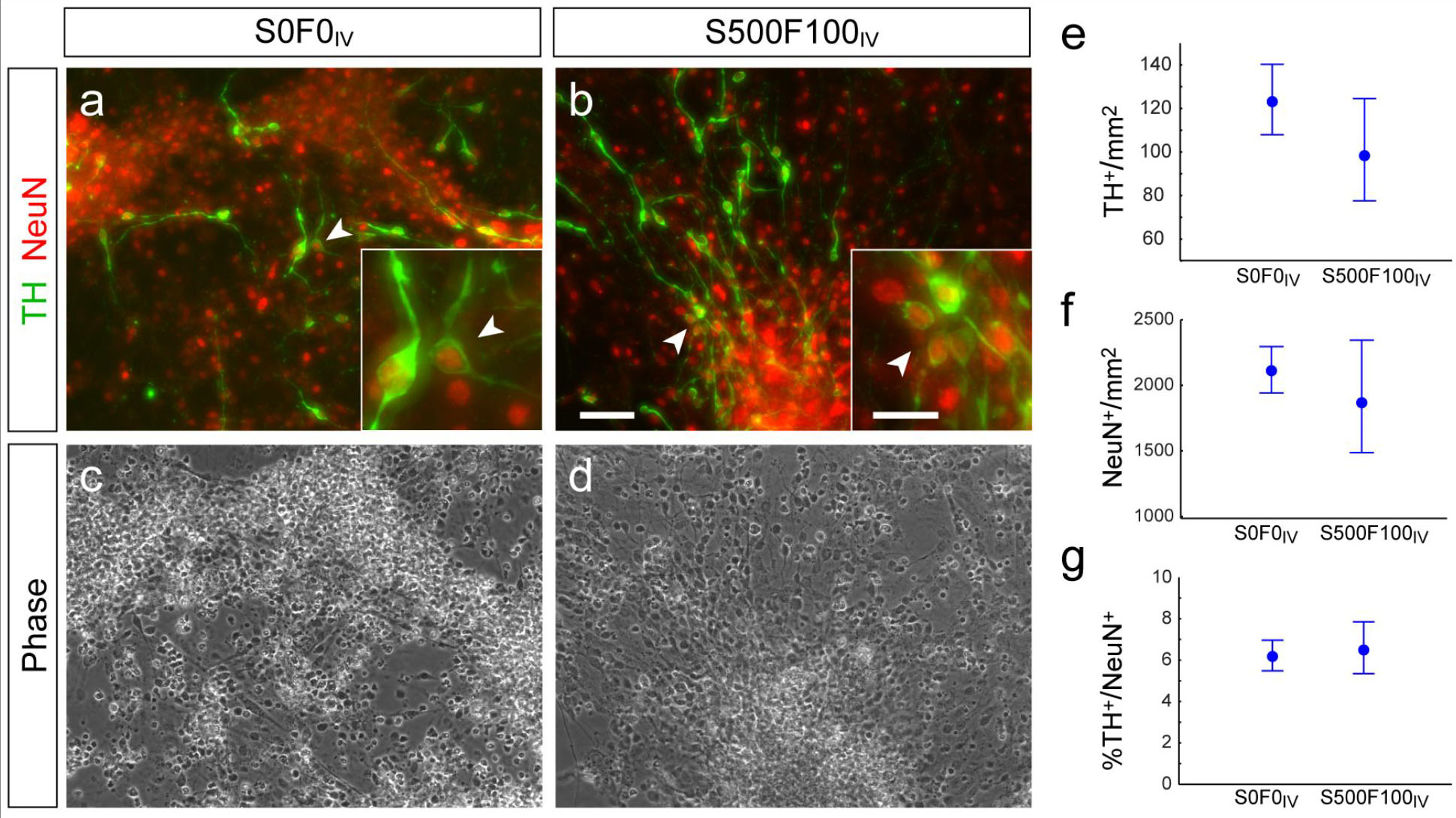
Exogenous SHH and FGF8 during NP expansion did not enhance DA neuron production. Exogenous SHH and FGF8 during NP expansion did not enhance DA neuron production. Neurons and DA neurons in S0F0_IV_ and in S500F100_IV_ were immunolabeled for NeuN and TH, respectively. (a, b) Images of TH (green) and NeuN (red) immunofluorescence in S0F0_IV_ (a) and S500F100_IV_ (b). Arrowheads show examples of double-positive neurons that were TH+NeuN+; insets show high magnification images. (c, d) Corresponding phase contrast images. (e-g) Quantification of TH+ (e) and NeuN+ neurons (f) per unit area and percent of TH+ neurons versus NeuN+ neurons (g) showed no difference between S0F0_IV_ and S500F100_IV_ cultures. Means and 95% CI are shown Scale bar (a - d), 50 μm; insets, 20 μm. Values are in Table 2.

### The role of exogenous SHH and FGF8 on developmental signaling pathways

SHH, FGF8, and WNTs are maintained by interdependent and hierarchical genetic loops during embryogenesis that also function during dopamine neuron development *in vivo* [reviewed in 6, 35, 38]. Therefore, we determined whether endogenous SHH, FGF8, and WNT signaling pathway components in ESCs programmed to become DA neurons were affected by the addition of exogenous SHH and FGF8. We performed quantitative RT-PCR analysis on D3 ESCs and D3 ESC-derived neuronal precursors at the end of Stage IV in S0F0_IV_ and S500F100_IV_ conditions. We analyzed the mRNA expression of *Shh* and *Gli1* (a downstream transcription factor and high fidelity readout of the SHH signaling pathway) [39, 40]. We also assessed *Fgf8* and *Fgfr3*(which encodes a receptor required for FGF8 signaling in DA neurons) [41–45]. As shown in **Figure 5a**, *Shh*expression was nearly undetectable in Stage I ESCs but was highly expressed at Stage IV in both S0F0_IV_ and S500F100_IV_. However, exogenous SHH and FGF8 in S500F100_IV_ did not increase *Shh*expression (6296.52 ± 2014.11, cycle threshold) compared to S0F0_IV_ baseline (6184.75 ± 551.23, p = 0.9306). *Gli1* was detected in Stage I ESCs and was upregulated at the end of Stage IV. However, there was no difference in *Gli1* in the presence of exogenous morphogens (5.90 ± 2.39) compared to S0F0_IV_ baseline (5.89 ± 1.13, p = 0.9961) (**Figure 5b**). *Fgf8* and *Fgfr3* were also expressed in Stage I ESCs and upregulated at Stage IV. Treatment with exogenous morphogens did not alter *Fgf8* expression (3.68 ± 0.27) compared to the S0F0_IV_ (3.38 ± 0.53, p = 0.4253) (**Figure 5c**). *Fgf3r* levels in S500F100_IV_ (12.59 ± 3.40) were lower compared to the S0F0_IV_ (23.87 ± 4.00, p = 0.0205) (**Figure 5d**). Finally, *Wnt1* as well as *Nfat* and *Tcf4* (downstream components in WNT signaling) were expressed at low levels in Stage I ESCs and up-regulated at Stage IV (**Figure 5e, f**). Again, exogenous morphogens failed to alter the expression of these genes: *Wnt1* levels in the presence of morphogens was 175.80 ± 37.10 and without was 140.37 ± 48.24, p = 0.3703, **Figure 5e**); *Tcf4* in the presence of morphogens was 5.29 ± 1.17 and without was 6.49 ± 0.86 (**Figure 5f**, p = 0.2268); *Nfat* in the presence of morphogens was 2.87 ± 0.01 and without was 2.38 ± 0.70 (p = 0.2268). The expression levels of these transcripts in ESC-derived NPs are in good agreement with qRT-PCR analysis of DA neuron progenitors isolated *in vivo* [42].

**Figure 5.**
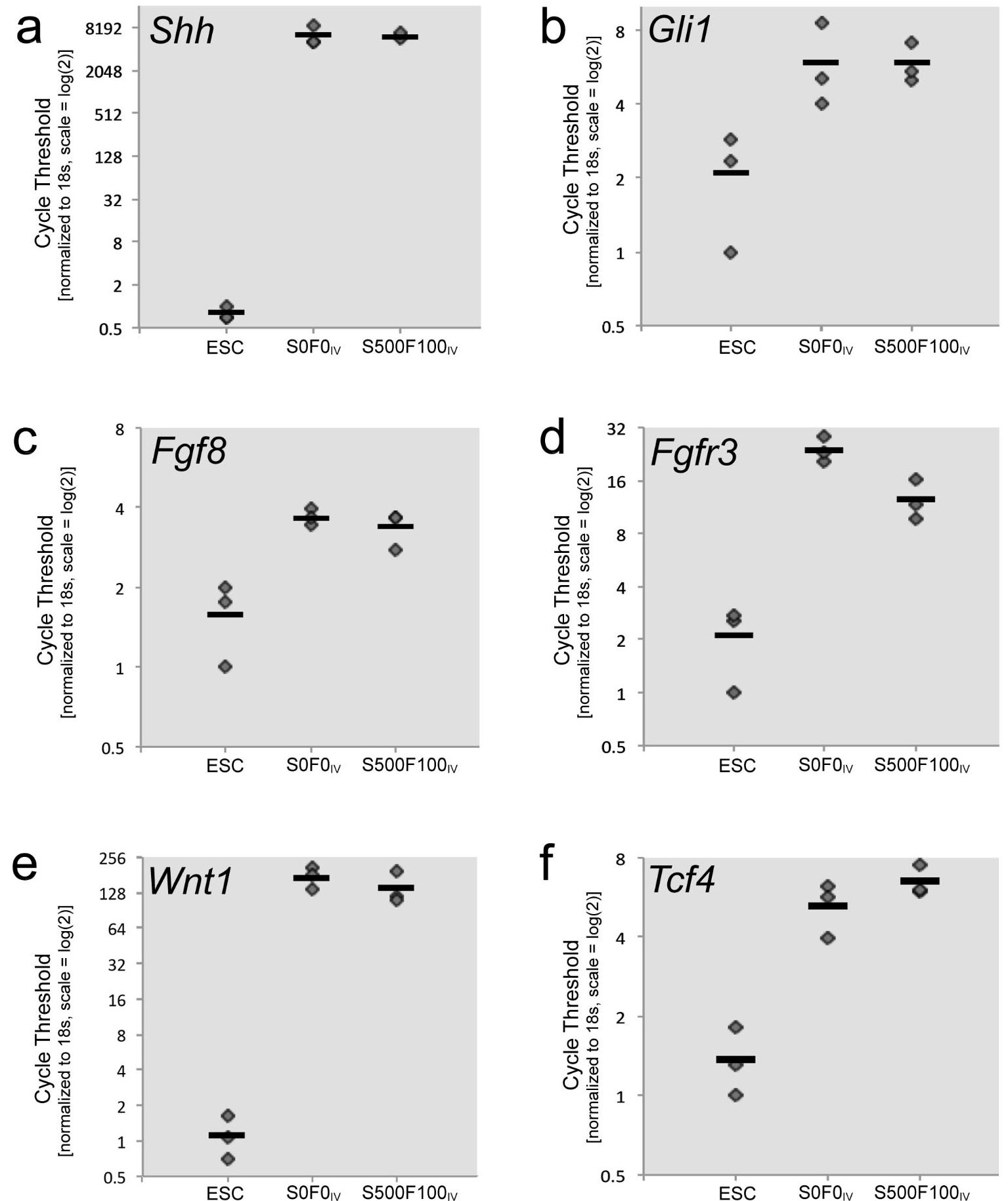
Endogenous SHH, FGF8, and WNT1 signaling components in ESCs and NPs. RNA was harvested from NPs at the end of Stage IV and compared to ESCs by qRT-PCR analysis. Expression of *Shh***(a)**, *Gli1***(b)**, which is a high fidelity readout of SHH signaling, and *Fgf8***(c)** increased during the transition from ESC to NP, but these genes were not affected by the exogenous morphogens SHH and FGF8. In contrast, *Fgfr3***(d)**, which encodes a receptor for FGF8 was decreased in the presence of exogenous SHH and FGF8. *Wnt1***(e)** and *Tcf4***(f)**, which is activated by canonical WNT signaling were increased in NPs compared to ESCs but unaffected by exogenous morphogens. The Y-axis is in log scale 2 to match cycle fractions. Three biological replicates are indicated by diamonds (and sometimes obscured by close values of replicates); the bar indicates the mean. The values and standard deviations are indicated in the results section.

### Comparative analysis with R1 ESCs: DA neuron production and subtype generation

To investigate whether the results we observed were unique to the D3 ESCs, we performed the same experiments using R1 mouse ESCs. R1 ESCs were cultured using the same baseline controland experimental conditions (S0F0_IV_ vs. S500F100_IV_). TH+ neuron production in S0F0_IV_ (186.65 TH+/mm^2^) was not significantly different from S500F100_IV_ (202.12 TH+/mm^2^) (p = 0.508, **Table 2** and **Figure 6**). These results show that similar to D3 ESCs, the production of TH+ neurons from R1 ESCs was not enhanced by the presence of exogenous SHH and FGF8 during the NP expansion stage. R1 ESCs also produced both CALB+ and CALR+ subtypes of DA neurons. The CALB+subtype (10.92%) was significantly higher than the CALR+ subtype (2.41%) (p < 0.0001) in S0F0_IV_ (**Figure 6f** and **Table 1**). In the presence of exogenous SHH and FGF8 in the S500F100_IV_ experimental condition, CALB+ DA neurons (10.87%) were significantly higher than CALR+ DA neurons (3.49%, p < 0.0001). The absence or presence of SHH and FGF8 did not result in any difference in the specification of CALB+ (p = 1.000) or CALR+ subtypes (p = 0.144). These results showed that R1 ESCs yielded more of the CALB+ subtype of DA neurons over the CALR+ subtype and that exogenous SHH and FGF8 did not shift this distribution - similar to findings with D3 ESCs.

**Figure 6.**
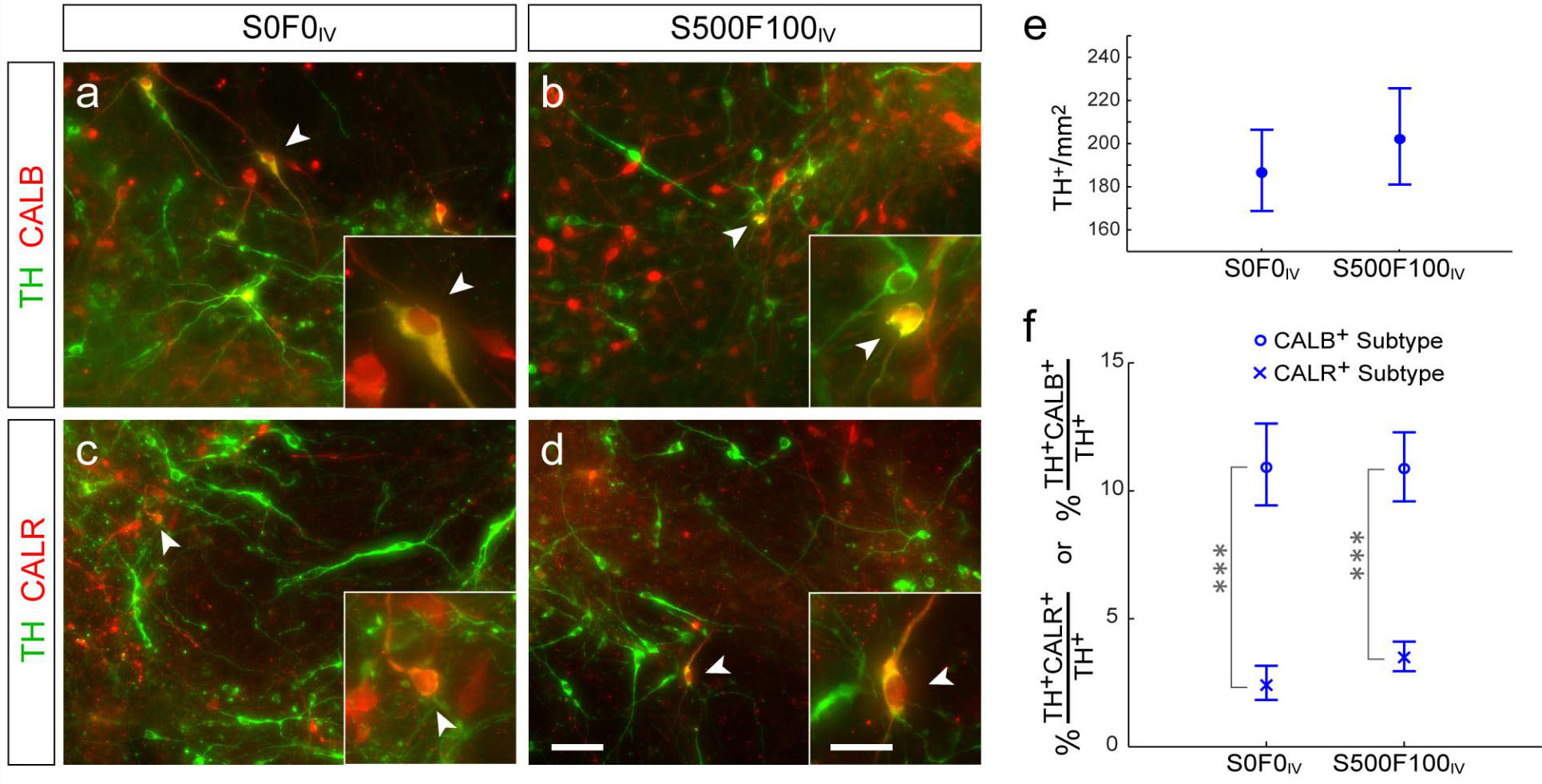
CALB+ and CALR+ subtypes of DA neurons were generated from R1 ESCs. The S0F0_IV_and the S500F100_IV_ conditions were repeated using R1 ESCs. **(a, b)** TH (green) and CALB (red) immunofluorescent labeling in S0F0_IV_ **(a)** and S500F100_IV_ **(b)**. **(c, d)** Images of TH (green) and CALR (red) immunofluorescent labeling in S0F0_IV_ **(c)** and S500F100_IV_ **(d)**. Arrowheads indicate examples of neurons that are double-positive for TH+CALB+ or TH+CALR+, and high magnification images of the same region are shown in the insets. **(e)** Quantification of TH+ neurons showed no difference in DA neuron production with the addition of SHH and FGF8. **(f)** Quantification of CALB+ and CALR+ subtypes of DA neurons revealed a higher percent of CALB+ than CALR+ subtype in both S0F0_IV_ and S500F100_IV_ cultures, but no difference in CALB+ or CALR+ subtype when compared across culture conditions. Means and 95% CI are shown with values provided in Tables 1 and 2. *** p < 0.0001. Scale bar **(a-d)**, 50 μm; insets, 20 μm.

### Exogenous SHH and FGF8 during NP selection (Stage III) did not alter the proportion of DA neuron subtypes derived from D3 ESCs

Previous studies suggested that SHH and FGF8 did not affect DA neuron production when added before Stage IV [30]. Immunolabeling of ESC-derived NPs at Stage III revealed that LMX1a, a transcription factor and determinant of DA neurons, was distributed amongst Nestin+ NPs (**Supplemental Figure 4**). We therefore tested whether exposure to exogenous SHH and FGF8 at this earlier stage (Stage III) impacted DA neuron subtype generation. As illustrated in **Figure 7a-d**, SHH (500 ng/ml) and FGF8 (100 ng/mL) or no morphogens were added to the culture only during Stage III (denoted as S500F100III) (**Figure 7b, d, e**). Paired controls were cultured using baseline culture conditions (denoted as S0F0_III_) (**Figure 7a, c, e**). CALB+ (12.31%) DA neurons were more abundant than the CALR+ subtype (8.53%, p < 0.001) in S0F0_III_ (**Figure 7e**, detailed in **Table 1
**).

**Figure 7.**
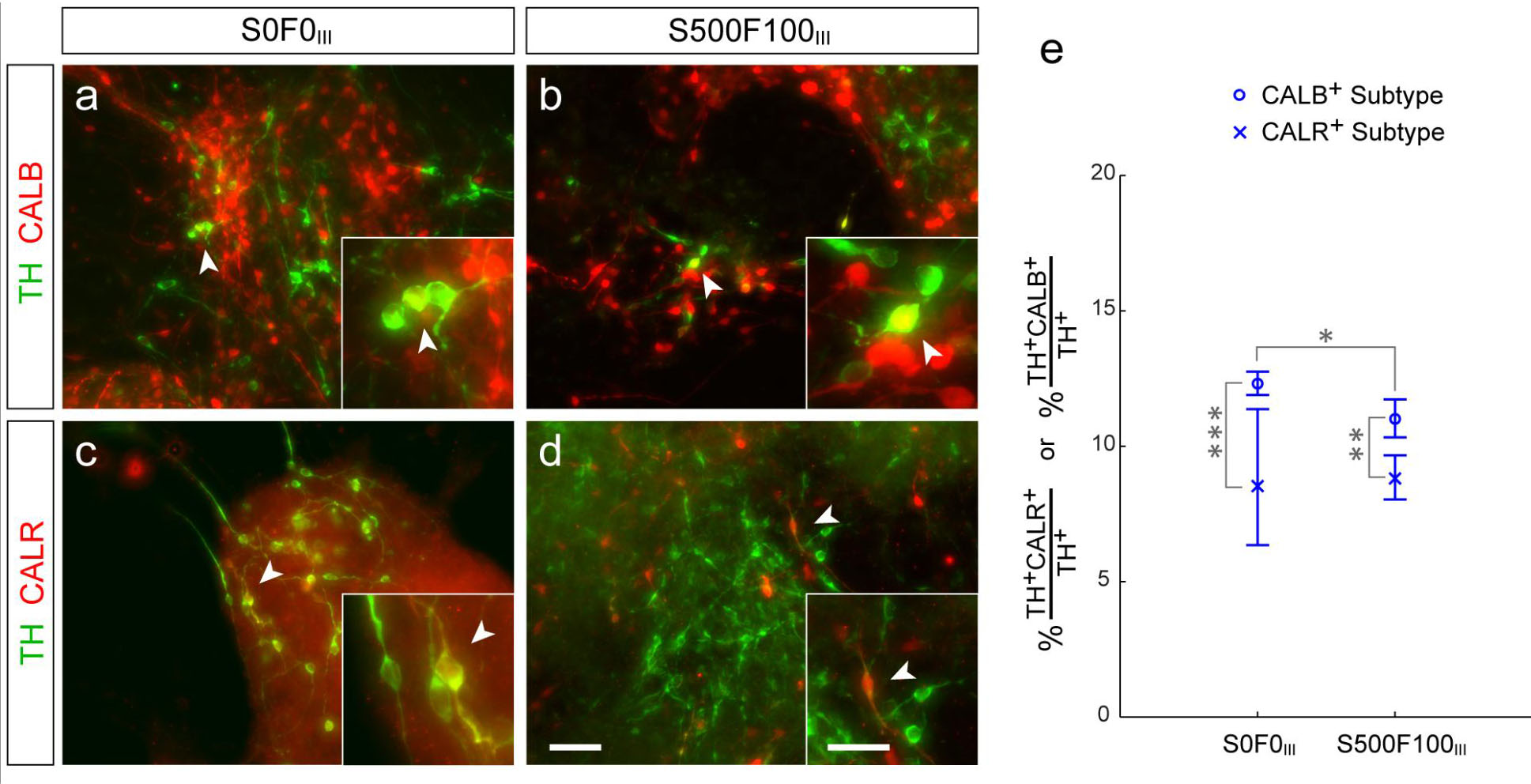
SHH and FGF8 at Stage III did not substantially govern CALB+ DA neuron subtype generation. D3 ESCs were cultured in parallel in the S0F0_III_baseline condition and in the S500F100_III_ condition, in which exogenous SHH (500 ng/ml) and FGF8 (100 ng/ml) were added during NP selection (Stage III) (a, b) Images of TH (green) and CALB (red) immunofluorescent labeling in S0F0_III_ **(a)** and S500F100III **(b)**. **(c, d)** Images of TH (green) and CALR (red) immunofluorescent labeling in S0F0_III_ **(c)** and S500F100III **(d)** conditions. Arrowheads indicate examples of neurons that were double-positive for TH+CALB+ or TH+CALR+; high magnification images of the same region are shown in insets. **(e)** Quantification of CALB+ subtypes and CALR+ subtypes of DA neurons revealed higher percent of CALB+ than CALR+ subtype in both S0F0_III_ and S500F100III cultures. The addition of SHH and FGF8 resulted in a small, but statistically significant reduction in the CALB+ subtype. Means and 95% CI are shown with values provided in Table 1. * p = 0.025, ** p = 0.0142, *** p = 0.0003. Scale bar (a - d), 50 μm; insets, 20 μm.

The CALB+ subtype (11.01%) was also significantly higher than CALR+ subtype (8.81%, p = 0.0142) in S500F100_III_ (**Figure 7e**). Comparison of CALB+ subtypes of DA neurons across culture conditions revealed a 1.30% difference (p = 0.025). There was no difference in the CALR+ subtype between S0F0_III_ and S500F100III (p = 1.000). In addition, there was no increase in the production of neurons or TH+ neurons with the presence of SHH and FGF8 during NP selection. We did not observe differences in DA neuron production, neuron production, or the percentage of DA neurons to total neurons between S0F0_III_ and S500F100_III_ (**Supplemental Figure 5
** and **
Table 2
**).

### Treatment with SHH-and FGFR3-neutralizing antibodies did not alter CALB+ or CALR+ subtype generation

Because *Shh* and *Fgf8* expression were detected at Stage IV regardless of exogenous SHH and FGF8 (**Figure 5**), we used neutralizing antibodies to block endogenous SHH and FGFR3 (**Figure 8**); FGFR3 is the receptor for FGF8 [34]. Cells were incubated with anti-SHH (10 µg/mL) and anti-FGFR3 (10 µg/mL) antibodies from the beginning of Stage III through the end of Stage V (**Figure 1c**; denoted as αSαFr). Under these conditions, the cultures did not receive exogenous SHH and FGF8 (**Figure 8** and **Table 1**). The paired control group received neither exogenous SHH and FGF8, nor anti-SHH and anti-FGFR3, denoted as S0F0_α_. In S0F0_α_, the CALB+ subtype (12.49%) was significantly higher than the CALR+ subtype (9.04%, p < 0.0001), in agreement with the previous control groups. In αSαFr, the CALB+ subtype (12.43%) was higher than the CALR+ subtype (8.21%, p = 0.0084). There was no difference in the CALB+ (p = 1.000) or CALR+ subtype (p = 0.467) in S0F0_α_ and αSαFr conditions. We did not observe differences in DA neuron production or the percentage of DA neurons in S0F0_α_ and αSαFr conditions (**Supplemental Figure 5** and **Table 2**).

**Figure 8.**
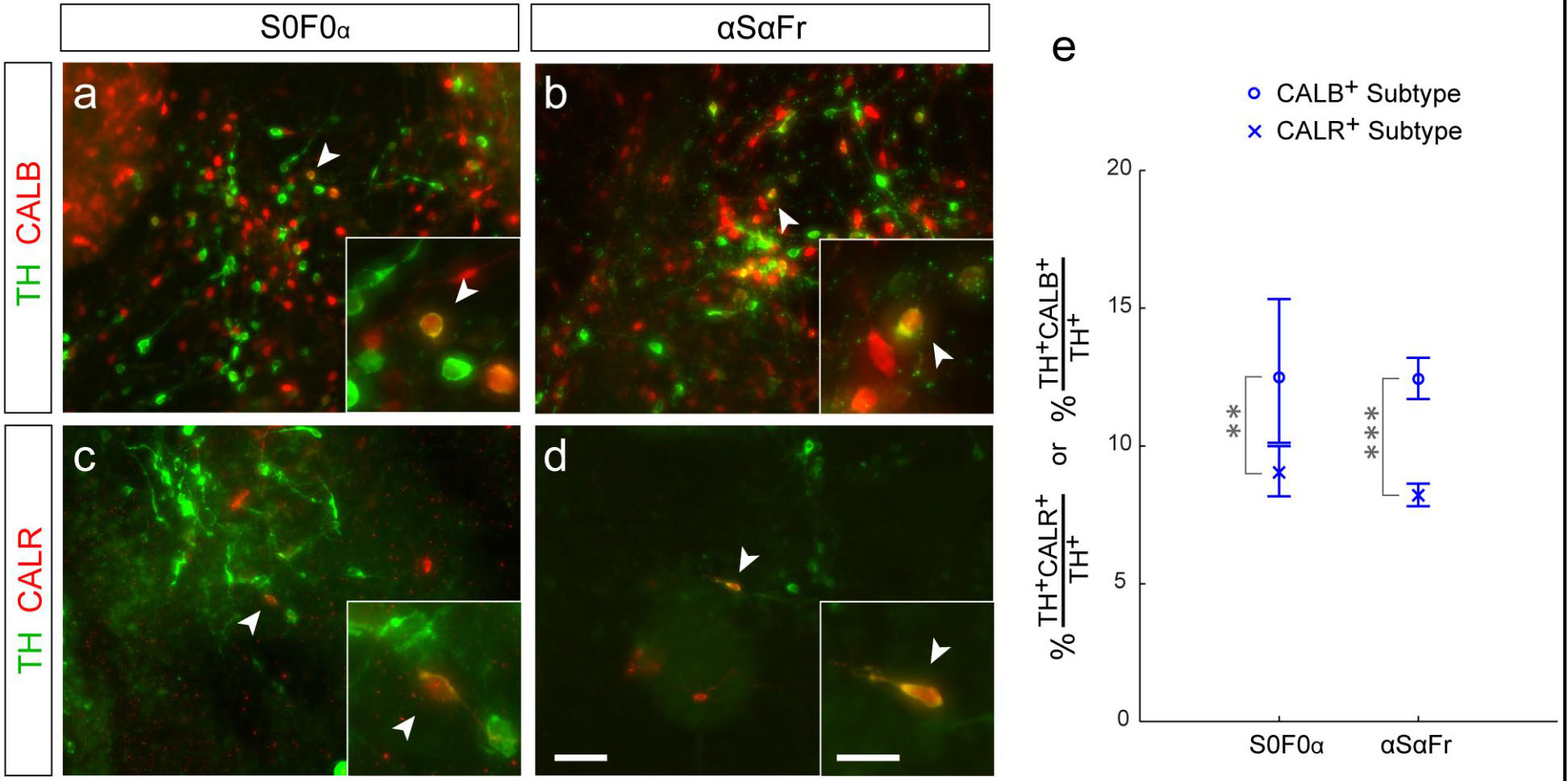
Blocking endogenous SHH and FGF8 signaling did not alter DA neuron subtypes. Blocking endogenous SHH and FGF8 signaling did not alter DA neuron subtypes. D3 ESCs were cultured in parallel in the S0F0_α_ baseline condition or in the αSαFr condition, in which anti-SHH (10 µg/mL) and anti-FGFR3 (10 µg/mL) neutralizing antibodies were added from Stage III through Stage V. (a, b) Images of TH (green) and CALB (red) immunofluorescent labeling in S0F0_α_ (a) and αSαFr (b). (c, d) Images of TH (green) and CALR (red) immunofluorescent labeling in S0F0_α_ (c) and αSαFr (d). Arrowheads indicate examples of neurons that were double-positive for TH+CALB+ or TH+CALR+, and high magnification images of the same region are shown in the insets. (e) Quantification of CALB+ subtypes and CALR+ subtypes of DA neurons revealed higher percent CALB+ than CALR+ subtype in both S0F0_α_ and αSαFr cultures. The addition of anti-SHH and anti-FGFR3 did not affect CALB+ and CALR+ subtype generation. Means and 95% CI are shown with values provided in Table 1. ** p = 0.0084, *** p < 0.0001. Scale bar (a - d), 50 μm; insets, 20 μm.

## DISCUSSION

Midbrain DA neurons *in vivo* consist of several subtypes with unique molecular identities and are involved in distinct neurological disorders [3,7,12]. An understanding of how specific subtypes of DA neurons are generated *in vitro* is important for developing appropriate research tools for studying disease mechanisms, investigating potential therapeutics, and adopting stem cells for clinical applications [8]. In this study, we differentiated DA neurons from ESCs using a 5-stage method and evaluated DA neuron subtypes by immunolabeling for CALB or CALR and TH and tested whether subtype generation *in vitro* was influenced by exogenous SHH and FGF8. Our initial observation of the differentiated cultures revealed a high degree of heterogeneity in the spatial distribution of all neurons and DA neurons. A brief survey of the literature reveals a limitation in the application and/or description of determining quantification methods for analyzing ESC-derived neurons. We developed a method to approximate errors associated with different sizes of random sampling, which was used to select the number of FOVs in our quantitative analysis. Our sampling and quantitative approaches can be applied to a variety of *in vitro* experiments as demonstrated here for assessing DA neuron diversity.

### The role of SHH and FGF8 in ESC-derived DA neurons

SHH and FGF8 are morphogens that function in embryonic patterning and play a role in the induction of DA neurons in the embryonic brain [34, 39–41]. These two morphogens have been suggested to enhance DA neuron production *in vitro* [30, 31, 46]. However, we did not observe an increase in the production of DA neurons in the presence of exogenous SHH and FGF8 from two independent ESC lines. There are several potential explanations for these findings including the following: 1) variations in ESCs and experimental procedures, 2) saturating levels of endogenous SHH and FGF8, and 3) context-dependent functions of SHH and FGF8.

The production of DA neurons from ESCs can be influenced by variations in cell line and experimental procedures. With regard to cell lines, we utilized both D3 and R1 ESCs for our experiments because these commercially available lines have been used previously in DA neuron differentiation experiments with the 5-stage protocol [30, 31, 47]. It is noteworthy that the descriptions of DA neuron production in the literature have indicated a range in the yield of DA neurons derived from ESCs treated with SHH and FGF8 (7% → 33% of total neurons reported by Lee et al. [30]; 3 →11% reported by Chung et al. [31]; 6% in this study). One experimental difference between previously published studies and ours is that we counted TH+ neurons versus NeuN+ neurons, which is restricted to neuronal nuclei and more amenable to cell counting than TUJ-1, which is expressed in neuronal soma and processes. However, it is also possible that our ESCs were at a passage that did not maximally respond to exogenous SHH and FGF8. Studies have reported variations in proliferation, differentiation, and genetic abnormalities with ESC subclones and passages [37, 48]. Because passage numbers of ESCs are often unknown or not reported in literature it is difficult to directly compare across studies, which necessitates the need to run untreated control groups. To eliminate reagent variability as a potential influence on cell responses, we compared reagents from a commercial DA neuron differentiation kit (R&D, sc001b) to reagents individually purchased (listed in Methods); we observed no differences in DA neuron production.

With regard to endogenous morphogen levels, our qRT-PCR analysis showed that mRNA levels of SHH and FGF8 signaling components were upregulated in cells at the NP expansion stage compared to undifferentiated ESCs indicating that some cells *in vitro* acquire robust expression of endogenous *Shh* and *Fgf8*. Along with the upregulation of *Wnt1*, these results are consistent with previous *in vitro* studies and are in alignment with *in vivo* development [30, 42]. The addition of exogenous SHH and FGF8 did not increase the expression levels of endogenous pathway components, but interestingly there was a decrease in the expression of *Fgfr3* in the presence of exogenous FGF8, consistent with a negative feedback loop of FGF8 signaling [49]. The dynamic change in intrinsic gene expression from ESCs to NPs and the lack of a robust functional response to exogenous SHH and FGF8 suggest that NPs produce sufficient and/or saturated levels endogenously and may be a self-sufficient system for limited DA neuron production. We directly tested the role of endogenous SHH and FGF8 signaling with neutralizing antibodies [34] and saw no decrease in the number of TH+ neurons. These results suggest that a low level of endogenous morphogen persisted in the presence of the antibodies or that a short pulse of morphogen activity occurred prior to blocking antibody disruption, which would be consistent with the dynamic temporal requirement for morphogen induction of DA neurons *in vivo*[14, 34, 41].

We confirmed the pluripotency of D3 ESCs with OCT4 expression, which was further supported by the ability to generate DA neurons, CALB+ and CALR+ neurons, and a substantial number of additional neurons. We also observed the acquisition of LMX1a expression in DA progenitors at an appropriate stage. However, our findings and previous studies collectively indicate that the majority (67 - 94%) of NPs do not differentiate into DA neurons in response to exogenous SHH or FGF8 [30, 31]. SHH and FGF8 have been observed to fail to transdifferentiate striatal neurons to express TH *in vitro* [50]. While SHH and FGF8 are potent morphogens for DA neuron induction *in vivo*, it is important to note that they act on specific precursor cells suggesting context-dependent functions of these morphogens. SHH is involved in the induction of not only DA neurons, but also serotoninergic neurons, motor neurons, and GABAergic interneurons. This shows that the same factor can have different effects depending on the recipient cells [36, 51–53]. Further, it is only within a critical window *in vivo* that SHH and FGF8 can induce ectopic DA neuron production [34]. It is interesting to note that when ESC-derived NPs overexpress key transcription factors such as *Nurr1* and *Lmx1a*, these NPs produced 60-80% DA neurons when exposed to SHH and FGF8 [31, 32, 47]. Taken together, these studies suggest that many of the microenvironmental cues that can influence cell fate may be missing *in vitro*. Alternatively, there may be an unidentified mechanism(s) that ensures only a limited number of progenitors are poised to acquire a DA neuron identity. Other mechanisms are likely to be involved with subtype specification, such as other soluble morphogens, progenitors niche, cell-cell communication, migration and refinement of progenitors, and intrinsic gene expression [54]. It is likely that additional intrinsic and extrinsic factors and mechanisms are required to work synergistically with exogenous SHH and FGF8 to enhance DA neuron production.

### DA neuron subtypes derived from mouse ESCs

After ten days of differentiation, a small percentage of DA neurons in our culture expressed CALB (11 - 27%) or CALR (2-13 %). Remarkably, our culture conditions recapitulated the relative distribution of these subtypes of DA neurons in the mouse brain, which also has a higher percentage of CALB+ versus CALR+ DA neurons in the midbrain [15, 16, 55]. The differences in percentages between DA neuron subtypes produced *in vitro* and established *in vivo* may reflect differences in the time, context, or embryonic niche where DA progenitors are located compared to culture conditions. CALB and CALR expression in DA neurons are not mutually exclusive *in vivo [16]*. Although we did not perform triple-immunolabeling of TH, CALB, and CALR due to antibody compatibility complications, it will be interesting in the future to quantify the co-expression of CALB and CALR in DA neurons. Because cognitive function and reward-associated behaviors involve DA neurons located in the VTA, which is high in CALB+ and CALR+ DA neuron subtypes, our findings regarding the *in vitro* production of these subtypes is potentially applicable to aid in the study of complex disorders such as schizophrenia, addiction, reward, and arousal. The GIRK2+ subtype of DA neurons in the SN is vulnerable to degeneration in Parkinson’s disease [12] and transplanting an enriched population of SN DA neurons has led to greater motor function recovery in a rat Parkinson’s disease model [4, 11]. In this study, we did not detect GIRK2 expression at the end of fairly standard differentiation duration (10-15 days) in the 5-stage paradigm. However, previous studies have detected GIRK2 expression in DA neurons derived from a similar paradigm after being engrafted in animal models with DA neuron lesions [32, 56, 57]. Thus, DA neurons derived from ESCs have the ability to become the GIRK2+ subtype of DA neurons. The absence of GIRK2 expression may be explained by the temporal dependency, intrinsic mechanisms, or microenvironment [39]. The assertion of temporal dependency is supported by our observation that a small number of GIRK2 expressing DA neurons was observed after 25 days *in vitro.*

Finally, we hypothesized that the generation of DA neuron subtypes is influenced by exogenous SHH and FGF8. The rationale for testing our hypothesis is that precursor cells in the developing midbrain are exposed to different concentrations of SHH and FGF8 secreted by the ventral midline and isthmus, respectively [34]. Subtype specification through a SHH gradient has been demonstrated in spinal cord motor neurons [36]. In addition, SHH and FGF8 gradients control cell fate specification in limb bud development [58, 59]. Surprisingly, the addition of SHH and FGF8 at the NP expansion stage did not alter the production of the CALB+ subtype of DA neuron and resulted in a minor decrease of CALR+ DA neurons. Even though the decrease was statistically significant, the change was very small and in our opinion unlikely to be an efficient method to manipulate DA neuron subtype generation *in vitro*. Similarly, neutralizing antibodies did not alter the number of CALB+ or CALR+ DA neuron subtypes. These findings suggest that subtype generation *in vitro* is not governed by exogenous SHH and FGF8, but may be controlled by intrinsic mechanisms such as transcription factors. We did note a significant difference in the percentages of both CALB+ and CALR+ DA neuron subtypes derived from earlier versus later ESC passage, although the pattern of more CALB+ than CALR+ DA neurons remained. These results suggest that DA subtype specification may be sensitive to intrinsic factors and starting ESC passage, which agrees with literature that reported variability in the yield of differentiated cells among subclones of the same ESCs [37].

## CONCLUSION

We established a rigorous analytical approach to sampling and quantify DA neuron subtypes generated from ESCs using a 5-stage method. Higher percentages of CALB+ than CALR+ subtype of DA neurons were produced with this method. Manipulating SHH and FGF8 signaling with exogenous SHH/FGF8 addition or SHH-and FGFR3-neutralizing antibodies was not effective in altering subtype generation *in vitro*. Taken together, our findings point toward endogenous mechanisms that govern DA neuron production and subtype differentiation of CALB+ and CALR+ DA neurons from mouse ESCs using the 5-stage method. Subtype specification likely involves multiple complex mechanisms that were not identified in this study and not controlled solely by SHH and FGF8. A deeper understanding of intrinsic gene expression patterns, both spatial and temporal, and their relation to allocating DA progenitors and subtype specification will be particularly useful for the field of ESC and DA neuron research.

## Acknowledgements

The authors thank Patrick Dingle for the Monte Carlo simulation Matlab script, Samantha Brady for assistance in quantification of R1 ESC studies, and Kirsten Sigrist and Marissa Furey at the Brown Transgenic Facility for sharing their knowledge of ESC techniques.

